# Microinductor-Fused Atomic Force Microscopy Cantilevers for Dynamic Imaging and Multimodal Manipulation

**DOI:** 10.1101/2021.05.13.444109

**Authors:** Charilaos Mousoulis, Xin Xu, Robert L. Wilson, Garrett Chado, Joseph Wahlquist, Mark P. Stoykovich, Virginia L. Ferguson, Babak Ziaie, Corey P. Neu

## Abstract

Recent advances in atomic force microscopy (AFM) imaging and force spectroscopy have demonstrated improvements in rapid acquisition of quantitative data for applications in materials science, surface characterization, and biology. However, conventional AFM technology is limited in detection sensitivity and the ability to excite at off-resonance frequencies restricting broad utility of the technology. Here we demonstrate new AFM cantilevers fabricated with a planar microcoil at the tip region, which can be used to generate or sense highly-localized magnetic fields. Torsion/bending actuation of the cantilevers is accomplished with simple experimental configurations, enabling quantitative and simultaneous mapping of both stiffness and friction at the sample surface with more than one order of magnitude improvement in compositional contrast. Our method is compatible with commercial AFM systems, allowing us to clearly resolve small stiffness and friction variations in copolymer and biological samples that were difficult to detect by conventional AFM methods. In combination with fluorescence microscopy, we also generated localized fields to selectively stimulate and monitor structural changes in viable cells with nm-scale detail. Hybrid AFM cantilevers may be useful to characterize a broad range of complex material surfaces, in addition to combined physical and chemical analyses of single cells and biological microenvironments.

## INTRODUCTION

AFM is a leading surface probing technology with diverse application in multiple fields in science, engineering, and medicine^1–6^. A significant advantage of AFM over electron or optical microscopy is that AFM provides information on the nanoscale mechanical and chemical properties of the sample surface^7–11^. While the utility of AFM is broad, there is still a need to improve the detection sensitivity of the method to enable detection of small surface variations and features, improve dynamic measurements of torsional motion, including friction and combined torsion/bending actuation, and probe and sense features below the surface that is typically not directly in contact with the cantilever. Consequently, in the past decade many new methods have been developed to improve AFM sensitivity for mapping material properties on nanoscale ^12–18^, including technologies that utilize modified AFM cantilevers^12,16–18^. On the other hand, AFM study of single cells has gained wide attention ^19–25^. AFM imaging works in cell culture medium without the need for cell fixation or staining; furthermore, AFM is also an essential tool to study cell mechanics, including stiffness, friction, adhesion, and morphology^26^.

We previously presented the concept of coupling AFM and nuclear magnetic resonance (NMR) technologies, through a modified commercial AFM cantilever, to enable highly-localized, cellular-scale nuclear magnetic resonance spectroscopy^27^. Utilizing hybrid cantilevers equipped with a planar coil of a few tens of micrometers in diameter, we were able to distinguish characteristic peaks of different solutions in a 500 MHz NMR magnet. The platform, however, was based on a commercial product that lacked a low spring constant and optimized electromagnetic design, with a parasitic capacitance that was detrimental to the sensing potential. Nevertheless, this accomplishment set the foundation for a technology that can obtain simultaneous topological, structural, mechanical and chemical profiling of cellular-scale biological samples.

Here we present a hybrid probe, designed to be effortlessly integrated with a commercial AFM instrument and with a sufficiently low spring constant (i.e. approximately 0.25 N/m) allowing for the study of soft biological samples (Figure 1; Supplemental Figures S1-S3). The implementation of a variety of designs for a planar coil (with one or more turns), at the free edge of cantilever (tip region), allows for the generation of highly localized magnetic fields, which not only opens the door for NMR on single cells^27^, but also improves the sensitivity of several AFM techniques by up to several orders of magnitude and enables manipulation of magnetic nanoparticles. Furthermore, due to its special design, this hybrid probe generates clean resonance peaks in liquids, which is essential for stable dynamic mode AFM imaging and quantitative AFM force spectroscopy. We also demonstrate that this probe can be easily excited in torsional vibration modes, which enables simultaneous stiffness and friction mapping of the sample surface. This new actuation method provides more than one order of magnitude improvement in sensitivity, thus was able to clearly resolve small stiffness and friction variations that were difficult to detect by conventional AFM methods. We then utilized these microinductor-enabled cantilevers to develop minimally-invasive techniques for the study of internal mechanical perturbation and mechanobiology measures. In combination with AFM and fluorescence microscopy detection capabilities, the localized field from the microcoils also enables the selective stimulation and monitoring of cells injected with superparamagnetic microbeads. Through the application of targeted magnetic forces, we were able to apply various waveforms to direct the microdisplacement of the injected beads to allow insight into the structural architecture of the cell. Coupling this method with AFM techniques provides further insight into internal and external cellular mechanics over time, and suggests broad potential applications.

**Figure 1.**
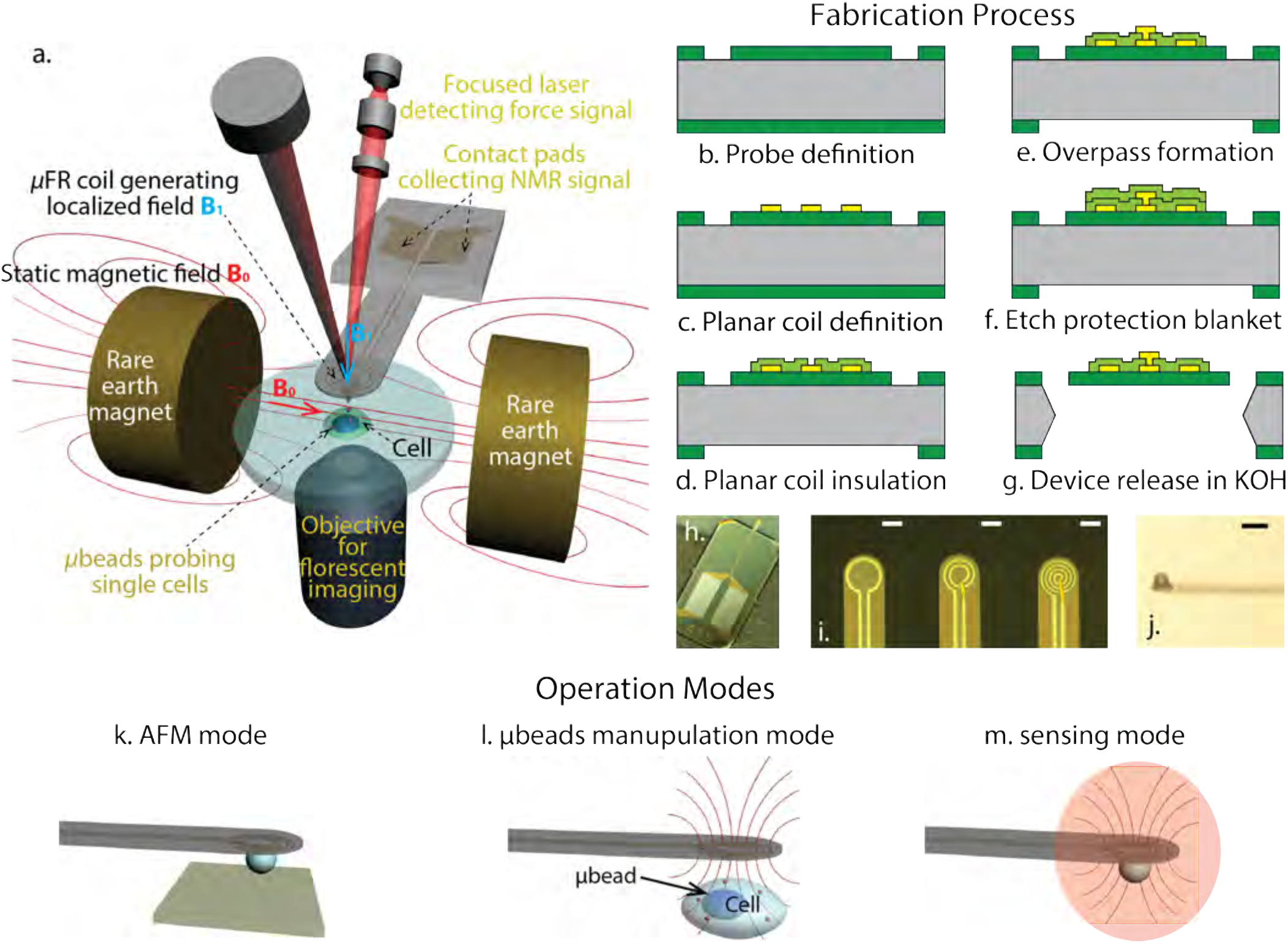
Concept of a multifunctional platform, fabrication process, and multiple operation modes of hybrid probe. (a) The concept of a multifunctional platform integrated with AFM, fluorescent microscopy and sensing (e.g. NMR) detection capabilities to sense and probe single cells using the hybrid probe. (b-g) Process flow for probes with multi-turn coils. (h) Optical microscopy photos of a 4-turn probe after its release. The different variations of the coil with 1, 2, and 4 turns are shown in (i). A side view of a cantilever after the attachment of the spherical tip is shown on (j). Operation modes of hybrid probe, including (k) AFM mode, where the transparency of the silicon substrate allows for exact positioning of the cantilever tip and the study of chemical functions via fluorescence microscopy; l) Microbeads manipulation mode, where the probe can be used to induce torque or displacements to embedded beads and track their location; (m) sensing mode, where the planar coil forms a highly localized region of magnetic field, therefore, when located in a homogeneous magnetic field environment, the planar coil enables nuclear magnetic resonance experiments for the study of chemical composition from picoliter-sized sample volume (i.e. at the scale of a single cell). Scale bars: 50 μm (i,j).

## FABRICATION AND VALIDATION

Microinductor-enabled cantilevers with various geometries were fabricated using microfabrication techniques in a well-defined, high-yield (60-70%) process (Figure 1(b-g), Supplemental Section 2). Briefly, the hybrid probe was comprised of a silicon nitride cantilever (300 μm long, 100 μm wide, 1 μm thick) on a silicon substrate. The traces of the planar coil, which were located at the free edge of the cantilever, extended to the base of the probe where they formed contact pads for the electrical connection to an external DC or AC circuit (Figure 1(h)). Three designs with varying coil turn quantities were implemented. The chosen numbers of turns for the planar coil were 1, 2, and 4, with corresponding average trace widths of 6.5, 6, and 1 μm approximately, and the spacing between the traces of the 2-turn and 4-turn coil was 6.5 and 9 μm respectively (Figure 1(i)). A glass spherical tip with average diameter of 20-27 μm (Cospheric LLC, Santa Barbara, CA) was manually attached to the edge of the cantilevers, with the application of UV-curable glue (NEA 121, Norland Products, Cranbury, NJ), for the purpose of AFM imaging (Figure 1(j)). The sensitivity and performance of the microcoil-enabled cantilevers improved with the number of turns, as evidenced by the corresponding increase in magnetic flux density observed near the center of the microcoils, enabling enhancements in probing by 1-2 orders of magnitude (Supplemental Figure S2). The magnetic field from the microcoils increases with input current, as expected, however at the cost of increased cantilever deflection and temperature. We simulated cantilever defection and temperature as a function of input current (Supplemental Section 4.5) and concluded that the changes in temperature and deflection do not affect AFM operation when the input current is < 1 mA.

## APPLICATIONS IN AFM IMAGING TECHNIQUES

Our microinductor-enabled cantilever can be used to enhance multiple AFM imaging modes (Figure 2). Piezo actuation is the most commonly used excitation method in dynamic mode AFM, wherein the cantilever is excited by high frequency vibrations from a piezoelectric transducer attached to the cantilever chip holder. However, it is well known that the piezo excitation in a liquid generates a “forest of peaks” in the cantilever response spectrum which are not necessarily related to the real cantilever resonances^28^. While some reported techniques address this issue^29–34^; our microinductor-enabled probe provides a simple and alternative approach. When current passes through the microcoil, the cantilever is locally heated and the thermal strain actuates the cantilever. Since only the cantilever is actuated, the cantilever response spectrum contains only the cantilever resonances even in liquids (Supplemental Figure S6), which is beneficial and essential for quantitative dynamic mode AFM spectroscopy in liquids^35^.

**Figure 2.**
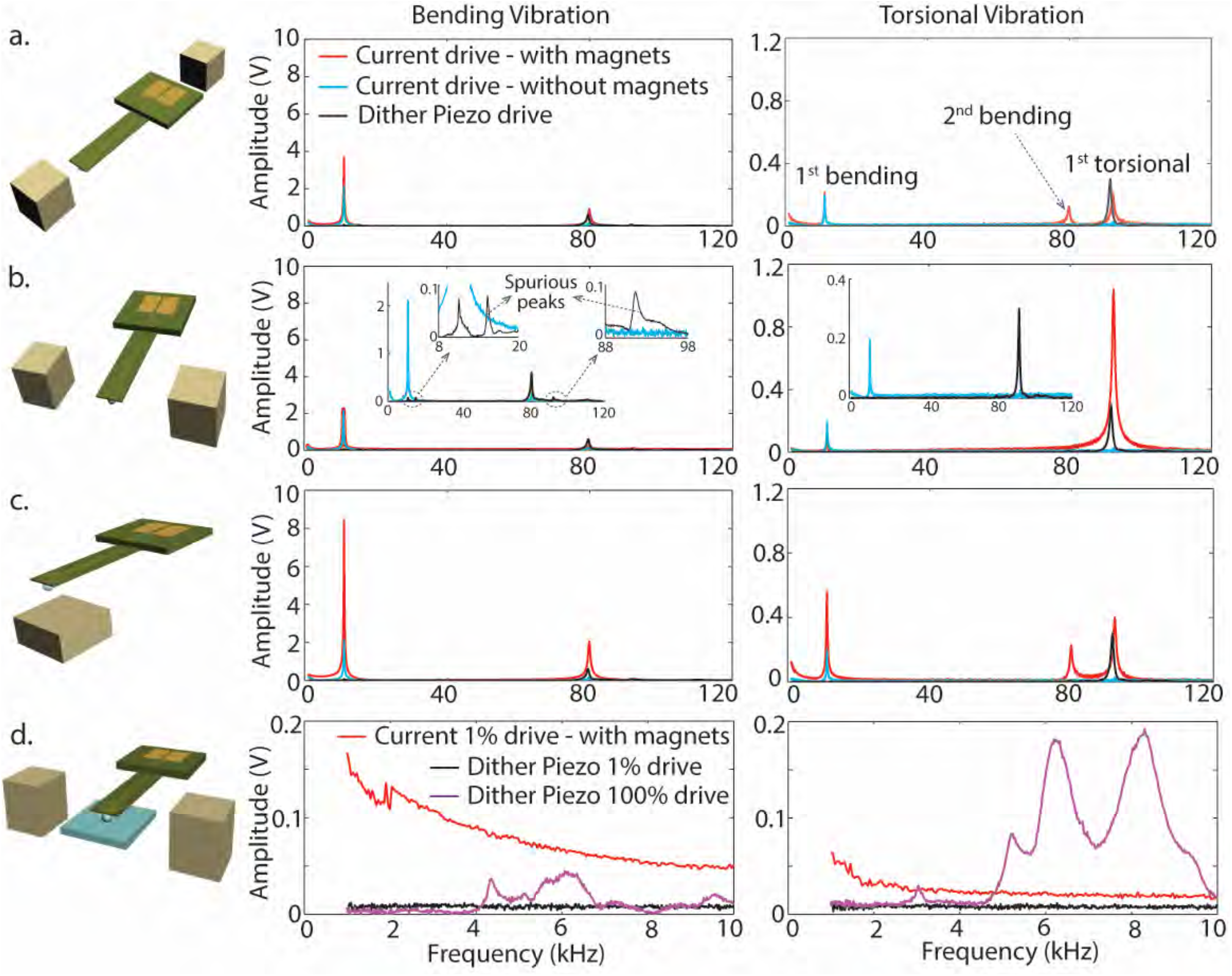
Simple magnet settings significantly enhance the bending and torsional vibration of coil probe. (a) Two small magnets placed at the longitudinal direction of the coil probe enhanced the 1^st^ bending mode free vibration by one order of magnitude. (b) Two small magnets placed transversely to the coil probe significantly enhanced the 1^st^ torsional mode free vibration. (c) A small magnet placed underneath the coil probe enhanced the 1^st^ bending mode free vibration by two orders of magnitude. (d) Two small magnets placed at the width direction of the coil probe enhanced both the bending and torsional modes vibration when the sphere tip is in contact with the sample (pre-load force 20nN). The inserts in (b) show the comparison of current drive without magnet enhancement (blue) and the dither piezo drive (black).

### Enhanced Bending and Torsional Modes

The AFM cantilever can be actuated in many eigenmodes of vibration each with its own natural frequency, including bending modes wherein the AFM tip is actuated normal to the sample surface, and torsional modes wherein the AFM tip is actuated parallel with respect to the sample surface. With simple configurations of permanent magnets around the coil probe, bending or torsional vibrations of the cantilever can be greatly enhanced. Experimental results of example configurations are summarized in Table 1. Larger drive amplitudes, stronger magnets, and smaller gaps between magnets further strengthen the magnetic field at the microcoil location and thus enhance the response amplitude by current drive. When the coil probe is placed between two 6.35 mm (¼”) cube neodymium magnets (Grade N52, K&J Magnetics Inc., Pipersville, PA) in the longitudinal direction (35 mm gap, ~10 mT magnetic flux density at the center), the 1^st^ bending mode vibration is enhanced 49 fold relative to the same cantilever excited by the piezo actuator (both are driven at 1% of the full drive range); even without the magnetic enhancement, the 1^st^ bending mode vibration is enhanced by 28 times using current drive (Figure 2(a)). Note that in the piezo drive spectrum, the 2^nd^ eigenmode vibration is enhanced; however, for such soft cantilevers (~0.25 N/m), the 1^st^ eigenmode vibration is more difficult to excite. Since the 1^st^ eigenmode stiffness is only 1/39.2 of the 2^nd^ eigenmode stiffness ^36^, the 1^st^ eigenmode vibration is preferred for imaging soft samples. One can always increase the piezo drive to gain sufficient amplitude of the 1^st^ eigenmode vibration. However, with increased piezo drive, more distortion is observed in the 1^st^ eigenmode resonant peak; even more detrimental to imaging, other peaks that are not related to the cantilever resonances appear in the spectrum as the piezo drive increases (see insert of Figure 2(b)), making selection of real cantilever resonances difficult. Another way to enhance the bending vibration is to place a magnet underneath or above the coil probe. With a 6.35×6.35×3.175 mm^3^ (¼”×¼”×⅛”) magnet placed 500 μm below the coil (Figure 2(c)), the 1^st^ bending vibration is enhanced 2 orders of magnitude. Note that the relative position between the coil probe and the magnet will greatly affect the enhancement, the closer to the center of the magnet, the greater the enhancement (Supplemental Figure S7).

**Table 1.**
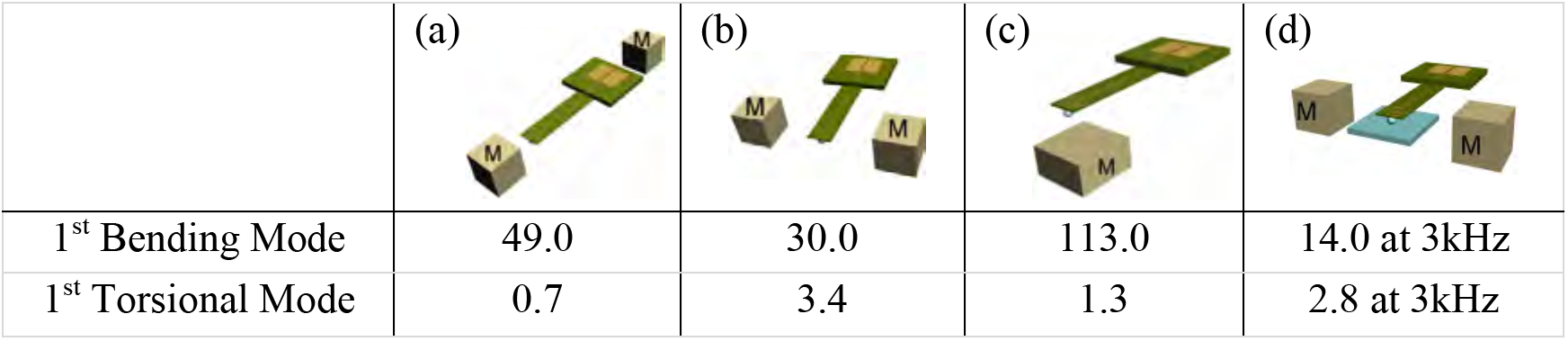
Magnet enhancement of current-driven bending and torsional vibrations relative to piezo drive. Vibration magnitudes were compared for the ratio of signal from current drive to piezo drive (factor improvement), both stimulated at 1% of the full drive range. For current drive without magnet enhancement, 1^st^ bending mode vibration is 28 times greater than that of piezo drive, 1^st^ torsional mode vibration is decreased to 0.08 times of that of piezo drive. To enhance bending vibrations, the magnets (marked with “M”) should be placed along the longitudinal direction of the cantilever as in mode (a) or below/above the coils as in mode (c); to enhance torsional vibrations, the magnets should be placed transversely to the cantilever as in mode (b) free vibration and (d) when in contact with sample.

|  | (a) | (b) | (c) | (d) |
| --- | --- | --- | --- | --- |
| 1<sup>st</sup> Bending Mode | 49.0 | 30.0 | 113.0 | 14.0 at 3kHz |
| 1<sup>st</sup> Torsional Mode | 0.7 | 3.4 | 1.3 | 2.8 at 3kHz |

To improve the torsional vibration, the coil probe can be placed between two 6.35 mm cube magnets in the width direction with 35mm gap (~10 mT at the coil probe location, Figure 2(b)). With this setting, the 1^st^ bending and 1^st^ torsional vibrations are enhanced 30 and 3.4 times as compared with the piezo excitation, respectively (both are driven at 1% of the full drive range). When the cantilever is in contact with a glass coverslip (with a preload force of ~20 nN; Figure 2(d)), it is difficult for the piezo actuator to excite bending mode vibrations in the low frequency range (working frequency range for force modulation mode) even with the maximum drive, whereas the current drive at 1% full drive range can easily enhance the bending vibration by one order of magnitude. To excite torsional vibrations when the cantilever is in contact with a sample using the piezo actuator, a large drive amplitude is needed. However, multiple peaks which are not all related to the cantilever resonances are shown in the response spectrum, making quantitative material property measurements impossible^37^. In comparison, current drive mode with only 1% drive yields a relative flat response for the torsional vibration and provides sufficient amplitude for force modulation mode imaging.

A significant advantage of the coil probe is that it can be *directly* excited for torsional vibrations in a simple experimental setup, even when in contact with a sample and off resonance (Figure 2, Table 1). The traditional excitation method for torsional vibration is to use a piezo actuator; however, this method introduces spurious resonances. A previous study proposed a method to directly actuate torsional vibration on the cantilever by attaching a micrometer-scale magnetic bead on the back side of the cantilever^17^. One challenge with this previous work is that the microbead needs to be carefully magnetized along the width of the cantilever, so that the cantilever can be torsionally actuated by a magnetic field produced from a solenoid placed underneath the sample. Although this method was able to efficiently excite torsional vibrations, small misalignments of the bead’s magnetic moment could introduce large distortions in the cantilever response. In contrast to this previous work, our microinductor-enhanced cantilever can be easily excited in the torsional vibration mode with a simple experimental setup.

### Improved Sensitivity and Contrast to Stiffness and Friction in Heterogeneous Systems

The application of the coil probe in bending (flexural) and torsional force modulation imaging is first demonstrated on a sample of 10 μm polystyrene (PS) beads (Life Technologies, Grand Island, NY, USA) embedded in polydimethylsiloxane (PDMS) elastomers (Sylgard 184 Silicone Elastomer, Dow Corning, Auburn, MI, USA) (Figure 3; Supplemental Figure S8). Force modulation AFM is widely used for mapping the local, nanoscale material properties by superimposing a low, fixed-frequency excitation on the cantilever in a constant force, contact mode scan^38^; the superimposed bending vibration amplitude is related to the compressive modulus of the sample, and the lateral vibration amplitude is related to surface friction and the lateral stiffness of the sample (Figure 3 top row). Since direct excitation is applied on the coil cantilever, the larger the vibration amplitude, the smaller the contact forces^37^. In this case, the compressive modulus of PDMS is 2-7 MPa^39^, while the compressive modulus of the PS bead is about 2 GPa^40^. However, the PS beads are possibly covered by a thin layer of PDMS making the bead area softer, but still much stiffer than pure PDMS. Thus, the bending vibration amplitude was higher on the softer PDMS (Figure 3 left panel). However, the torsional vibration amplitude was higher on the stiffer PS beads. The dominant force in this case was the dynamic friction between the AFM tip and the sample surface, not the lateral stiffness of the sample surface. The friction force was smaller on the PS beads than on the PDMS (Figure 3 right panel). For comparison, roughly the same imaging area was scanned by a standard sharp tip AFM cantilever (Multi75Al-G, NanoAndMore USA Corp., Watsonville, CA, USA) with piezo excitation. Since the excitation is indirectly applied to the cantilever, the imaging contrast is inverted from that by direct current drive^37^, i.e., the larger the vibration amplitude, the larger the contact forces. The normal vibration amplitude was larger on the stiffer PS beads (Figure 3). However, the torsional vibration amplitude showed little contrast between the PS beads and PDMS matrix due to very low torsional signal despite the piezo actuator being driven at maximum amplitude. This was observed in measurements of the contrast-to-noise ratio (contrast difference between the PS beads and PDMS matrix normalized by the standard deviation on a smooth region of PDMS); the contrast of friction mapping by conventional AFM was barely above the noise – thus the lack of sensitivity. Also, the spurious resonances could give inconsistent results^37^. Note that in both bending and torsional mode force modulation imaging, the measured vibration amplitude of the current-driven coil probe is significantly higher than the piezo-driven regular probe even though the current drive uses a much smaller driving voltage (0.08 V) than the piezo drive (8 V), which means the current drive provides a higher signal-to-noise ratio as well, especially for the torsional mode.

**Figure 3.**
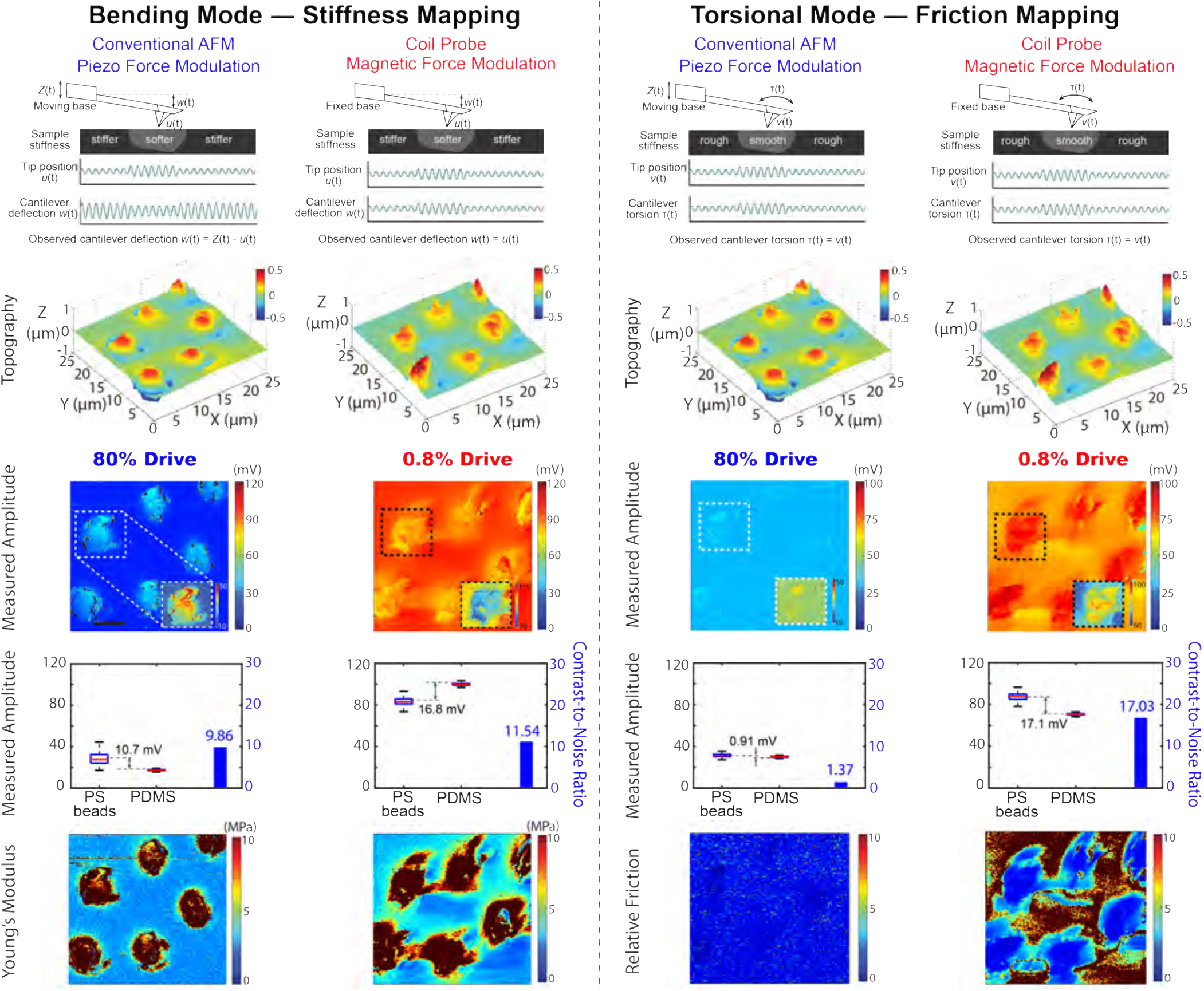
Current drive of the coil probe significantly enhanced the sensitivity of both stiffness and friction detection in force modulation mode imaging. The top row shows the schematics of bending and torsional modes of force modulation AFM using dither piezo and current excitation and corresponding waveforms (sample stiffness, cantilever deflection and tip position). Microbeads embedded in PDMS were scanned by regular probe with piezo drive, and coil probe with current drive. Note that although current drive uses a much smaller driving voltage (0.08 V) than the piezo drive (8 V), it yields a larger vibration amplitude, thus provides better signal-to-noise ratio. While both probes detected the stiffness variations, the coil probe is more efficient for friction mapping. Scale bars: 5 μm.

To further test the sensitivity of our technique, we compared the performance of the microinductor-enabled probe to the standard probe on substrates with subtle stiffness variations (Figure 4, Supplemental Section 6). First, we compared a scanned area of a PDMS sample that was covered by a thin layer of mica with an embedded microbead underneath (Figure 4(a)); the coil probe gave higher levels of detail on the stiffness variation due to the buried structures within the scan area as compared with the standard probe, demonstrating the higher sensitivity of the microinductor-enabled probe. Second, we scanned a 1×1 μm^2^ area of a PS-b-PMMA copolymer^41^ using the same coil probe operating in both current-driven and piezo-driven modes. Remarkably, even with a very blunt tip (~25 μm glass sphere), the current drive clearly resolved the small stiffness variations between the PS and PMMA domains in a lamellar pattern (Figure 4(b), Supplemental Figure S9) with a 47.5 nm periodicity (the bulk elastic moduli of PS and PMMA are 3.0 and 3.3 GPa, respectively^42^). In comparison, the piezo drive showed little contrast in the effective stiffness map even with maximum drive by the same blunt tip probe. To validate the nanostructures characterized by the current-driven operation using the coil probe, a standard sharp tip AFM cantilever was operated in force volume mode wherein an array of force-distance curves were collected over the entire scan area to generate a stiffness map^43^. Since the scan rate of the force volume mode imaging is very slow, we chose a smaller scan area of 200×200 nm^2^ with 64×64 pixels of resolution (Supplemental Figure S10). The same topography-stiffness relationship was found as in the force modulation imaging using current drive, *i.e.* the higher PMMA region is stiffer than the PS region^44^.

**Figure 4.**
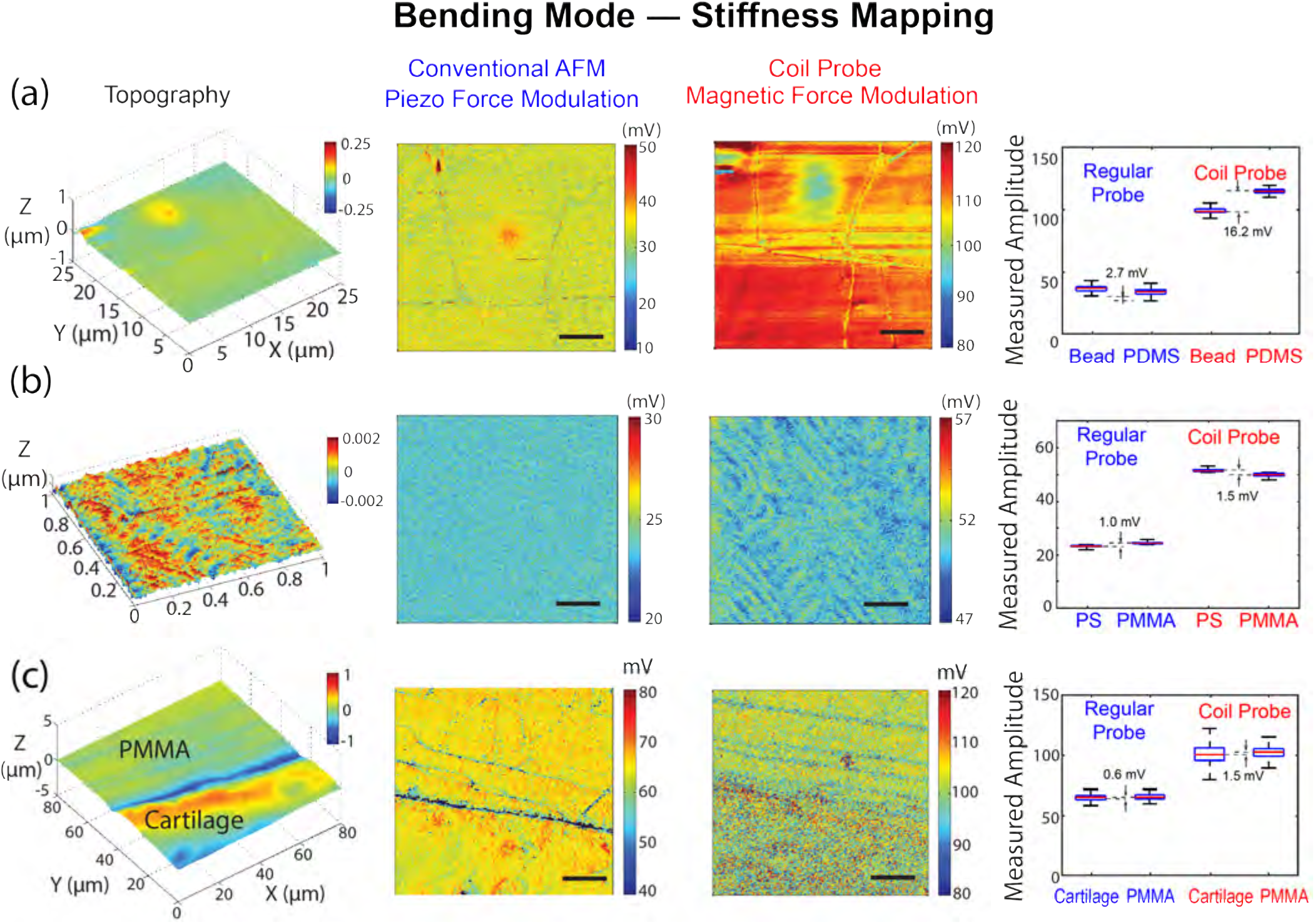
Current drive of the coil probe detects subtle stiffness variations that are difficult to detect by a conventional piezo drive with the same tip size. (a) A microbead embedded in PDMS covered by a thin layer of mica was scanned by conventional probe with piezo drive (i.e., piezo force modulation) and coil probe with current drive (i.e., magnetic force modulation), respectively; the coil probe provided more details on the stiffness map showing its high sensitivity. Scale bars: 5 μm. (b) Surface of a thin film of PS-b-PMMA copolymer scanned by coil probe with piezo drive and current drive respectively; while the current drive produced clear stiffness pattern, little contract was seen in the piezo drive map. Scale bars: 40 nm. (c) PMMA-cartilage sample was scanned by regular probe with piezo drive and coil probe with current drive respectively; even though the stiffness difference between the PMMA and PMMA penetrated cartilage is very small, the current drive is sensitive enough to detect the subtle difference. For all samples, current drive uses a much smaller driving voltage (0.08 V) than the piezo drive (8 V). Scale bars: 20 μm.

Finally, we further tested the coil probe on a PMMA-embedded articular cartilage sample (Figure 4(c), Supplemental Figure S11), and found that this technique was sensitive enough to detect the subtle differences in stiffness between PMMA and PMMA-embedded cartilage. Here, the coil probe was placed at the cartilage/PMMA interface region, which can be seen under the optical microscope. Since the PMMA is largely penetrated into the cartilage (~70-80% water volume fraction in the healthy tissue), the overall stiffness of the cartilage and PMMA regions are very close (<2% difference). However, from the effective stiffness map, these two regions can be easily distinguished: the PMMA region appears to be more homogeneous except the scratches due to polishing; while in the cartilage region, some surface zone locations are significantly softer than others. PMMA is a homogeneous polymer^45^, whereas PMMA embedded within the highly-anisotropic collagen matrix within demineralized cortical bone demonstrates a significant effect of orientation on nanomechanical properties^46^. Similarly, it is possible that varying orientation and prevalence of collagen fibrils and macromolecules within the PMMA-infilled articular cartilage contribute to variations in the effective stiffness and structural measurements ^47^.

We noted that dynamic friction mapping by torsional mode force modulation AFM also demonstrated enhanced signal amplitudes of approximately one order of magnitude compared to conventional AFM (Supplemental Figure S12). However, the samples selected (as mentioned: PDMS covered by a thin layer of mica, thin film of PS-b-PMMA copolymer, and PMMA-cartilage sample) all exhibited similar friction values and contrast when comparing piezo and magnetic force modulation due to the presence of polished or processed surfaces. Nevertheless, the enhanced amplitude of the measured signal for our hybrid probes indicates an efficient torsional actuation method for dynamic friction measurements and potentially lateral contract stiffness measurements in future applications.

## MANIPULATION AND ACTUATION OF MAGNETIC NANOPARTICLES

To demonstrate probing capabilities of the new microcoils, fluorescent superparamagnetic beads (0.9 μm) and carboxylate beads (1.1 μm) were ballistically injected into the cytoplasm or nucleus of bovine chondrocytes (Figure 5). Confocal microscopy (AIR_MP, Nikon) confirmed the penetration of the microbeads into the cytoplasm and nuclei. Microbead movements were accomplished using a 2- or 4-turn microcoil that was immersed in media and lowered to within 10-30 μm of the chondrocyte monolayer. A current pulse (0.2 mA, 0.5 s, 0.1 Hz) was passed to the cantilever. Cells were simultaneously imaged using widefield microscopy (Eclipse Ti, Nikon) and a CCD camera (iXon+, Andor, Belfast, Northern Ireland) for 60 s undisturbed, 120 s with pulsing, and a final 120 s undisturbed, at high spatiotemporal resolution (180 nm/pixel, 1.2 s frame rate).

**Figure 5.**
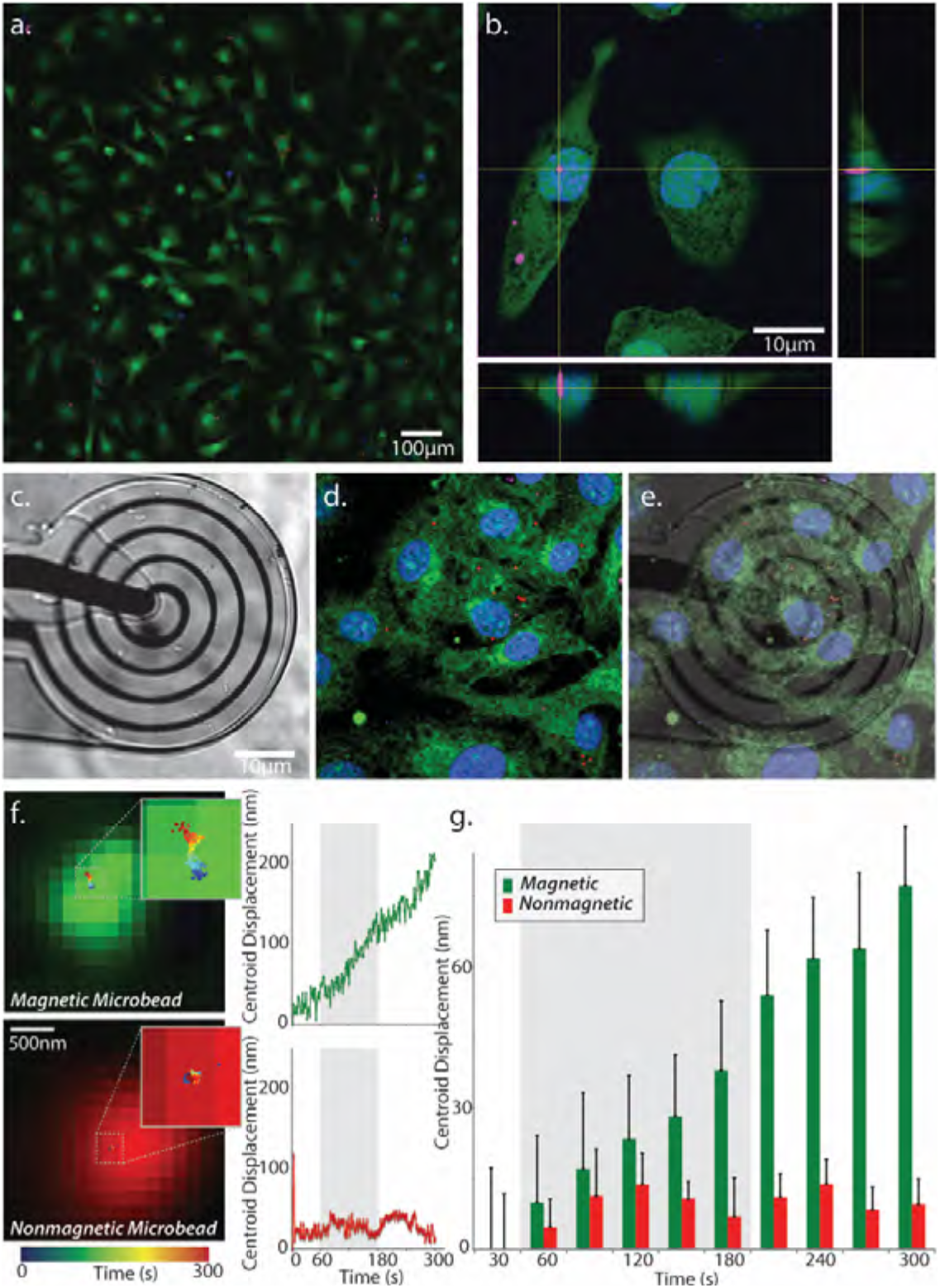
Localized fields from the hybrid probe displace superparamagnetic beads embedded within living cells. (a-b) Images (20×) of plated chondrocytes confirm microinjected superparamagnetic (magenta) and nonmagnetic (red) particles throughout the body and nucleus of cells. (c-e) A hybrid 4-turn microcoil was positioned 25 μm above the cells. (f-g) Cyclic magnetic fields (0.5 s application, 0.1 Hz) applied over 2 min induced nanometer scale displacements of the magnetic bead centroids, compared to controls.

Applied magnetic fields from hybrid microcoils resulted in motion of the superparamagnetic microbeads, but not for the nonmagnetic beads, as quantified by mean squared distances over time. A two-way ANOVA demonstrated significant effects of bead properties (i.e. magnetic particles versus nonmagnetic particles) and magnetic field application (p<0.01 for magnetic, p<0.01 for time, p<0.05 for interaction). Interestingly, the post-pulse section moved 14 nm more than the control of the same time period suggesting the possibility of a well-known creep response of the cytoplasm^48^. With an increase in temporal resolution, this creep behavior and the cellular microrheology may be modeled with this method^49^. It is important to note that heat production at an operating current of 0.2 mA does not increase the probe temperature beyond biologically relevant temperatures of 37 °C (Supplemental Figure S5).

Our microinductor-enabled cantilevers open the door for multiple new directions of research. First, the local probing of single cells in a population allows for decoupling of confounding factors (i.e. intracellular connections and signaling) present in the inherent setup of traditional magnetic tweezers by not applying the magnetic field to the entire testing dish, thus increasing testing efficiency and throughput. Second, the ability to perform localized measurements in media further enables the long-term testing of pharmacological agents designed to target rare (e.g. stem or cancer) cell populations. Third, coupled with visualization of localized nuclear architecture through chemical staining, it is possible to track changes in intranuclear biomechanics using deformable image registration to elucidate strain transfer from the cytoplasm and perinuclear regions to the nucleus interior ^50–52^. Additionally, the testing setup expands current capabilities of AFM through the simultaneous internal and external (surface) monitoring of mechanical and topographical measures, potentially revealing more information about this interconnection. Currently, improved insulation to allow for reliable electrical connections to the probe contact pads is needed during testing in fluid media, which is under further investigation. Nonuniform microparticle displacement within cells, evident in the large residual displacements, is also of concern and an aspect of current studies, potentially due to the heterogeneous actin network^48^.

## CONCLUSION

We have designed, fabricated and tested microinductor-enabled AFM cantilevers which can generate highly-localized magnetic fields. With a simple experimental setting, the bending and torsional vibration responses of the hybrid probe can be enhanced by at least one order of magnitude (sensitivity) compared with the most commonly used piezo excitation over a broader range of frequencies (bandwidth), thus has been utilized to enhance multiple AFM techniques, such as the tapping mode in liquids, as well as bending- and lateral-force modulation modes. Small stiffness and friction variations that were barely detected by conventional AFM methods have been clearly resolved by this new actuation method. Potentially, this technique can also be applied to improve the torsional resonance mode, torsional contact resonance mode, magnetic force microscopy, etc. We also demonstrated magnetic nanoparticles manipulation using this hybrid probe suggesting that this hybrid probe has tremendous potential for combined biophysical and biochemical analysis of single cells in complex *in vivo* microenvironments.

## ACKNOWLEDGEMENTS

This work was supported in part by NIH grants R01 AR063712, R21 AR066230, and R21 AR064178, and NSF grant CMMI CAREER 1349735. The authors thank Song Xu (Keysight Technologies Inc., Santa Rosa, CA, USA) for assistance in the design of the cantilever holder module, and for use of AC current supply modules.

## AUTHOR CONTRIBUTIONS

C.M., X.X., B.Z. and C.P.N. conceived of the study and designed the experiments. C.M. designed and fabricated the cantilevers and performed the finite element simulations. C.M., X.X. and R.L.W. performed the experiments described. G.C., J.W., M.P.S., and V.L.F. contributed essential reagents, samples and analysis. C.M., X.X., R.L.W., M.P.S. and C.P.N. wrote the manuscript. All authors reviewed the manuscript.

## SUPPLEMENTAL INFORMATION

### 1. Hybrid probe structure

**Figure S1.**
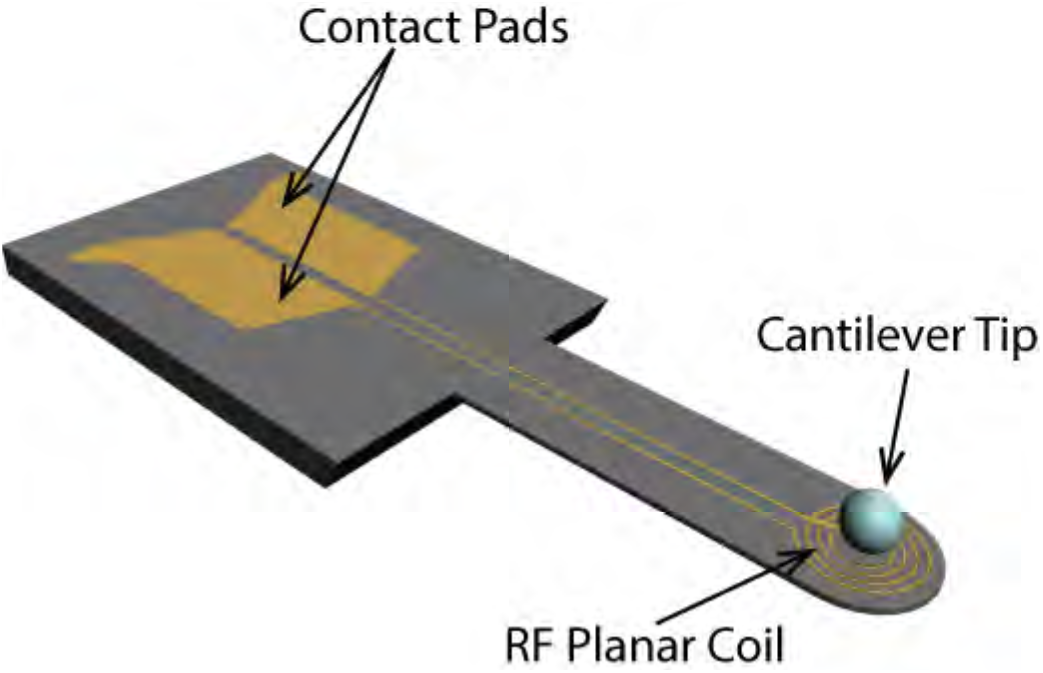
Hybrid probe structure. A silicon nitride cantilever, supported on silicon substrate, contains a planar coil of one or more turns that allows for electromagnetic experiments while the sample is scanned and information about its structure, stiffness, and adhesion to the substrate is collected.

### 2. Hybrid probe fabrication process

The fabrication process (Figure 1(b-g)) began with high resistivity (> 10,000 ohm-cm) 4 inch silicon wafers. The thickness of the wafers was approximately 300 microns. Both sides of the wafers had a 1 micron layer of low stress (tensile stress <100 MPa) low-pressure chemical vapor deposition (LPCVD) silicon nitride on both sides (Virginia Semiconductor Inc. Fredericksburg, Virginia). Initially, the nitride layer of the front of the wafers was patterned through lithography and SF_6_ dry etching. The bottom layer of the coils was then formed through electron-beam evaporation of 5 nm Cr (serving as an adhesion layer) and 25 nm Au. Application of approximately 100 nm plasma-enhanced chemical vapor deposition (PECVD) silicon nitride followed for the passivation of the metal layer. The deposition was split into three steps for the elimination of potential pinholes which would inhibit leakage currents and decrease etching resistance. The passivation layer was then patterned and etched in order to form the required vias for the electrical connection of the overpass to the covered trace of the multi-turn coil. The etching depth was controlled with high precision for the protection of the underlying cantilever. The back side LPCVD nitride was then similarly patterned through dry etching. The rest of the processing was carried out on the front side of the wafers. The top layer of Au (overpass) was deposited using electron-beam evaporation (10 nm Cr, 150 nm Au) and patterned using lift-off. An additional PECVD silicon nitride (approximately 340 nm) provided the final passivation of the top probe surface for the protection of the metal layers and the contacts from the KOH solution. Etching of the silicon substrate was then performed for approximately 3 hours in the KOH solution (45% w/v ratio in deionized H_2_O, 80 °C, with the addition of isopropyl alcohol). Finally, the excess PECVD nitride was removed through plasma etching.

Three designs with different coil turns were implemented. The chosen number of turns for the planar coil were 1, 2, and 4 with corresponding average trace widths of 6.5, 6, and 1 μm approximately, after fabrication (Figure 1(i)). Following the release of the devices and their detachment from the wafer, a glass spherical tip with average diameter of 20-27 μm (Cospheric LLC, Santa Barbara, CA) was attached to the edge of the cantilevers manually (Figure 1(j)), with the application of UV-curable glue (NEA 121, Norland Products, Cranbury, NJ). Using thermal noise analysis^1^, the resonant frequency was measured to be approximately 10 kHz with a spring constant of 0.25 N/m. These characteristics allow for the study of soft biological samples, e.g. tissues and cells, in air and liquid.

The fabrication process of the hybrid probes can be performed with high yield at any cleanroom equipped with the described MEMS manufacturing tools. However, the tip attachment process was manual and was performed under a microscope. To increase throughput and reduce labor, a semi-automatic solution should be explored for tip attachment, e.g. transfer of the adhesive agent and tip attachment using the positioning of an AFM system.

### 3. Magnetic Field Simulation

The sensitivity and performance of the microcoil-enabled cantilevers improved with the number of turns, as evidenced by the corresponding increase in magnetic flux density observed near the center of the microcoils, enabling enhancements in probing by 1-2 orders of magnitude (Figure S2, ANSYS Maxwell). Therefore, the performance of the hybrid probe depends strongly on the magnitude and spatial distribution of the magnetic field generated in the proximity of the cantilever tip. We simulated the magnitude of two orthogonal components of the magnetic flux density for a planar coil with 1, 2 and 4 turns in an area that extends up to 70 μm away from the center of the coil (point 0 of the horizontal axis) and up to 100 μm from its surface (point 0 of the vertical axis). The outer diameter of the coil in all cases was between 75 to 90 μm. The geometry of the coil allowed for the assumption of cylindrical symmetry, and therefore the two components *x* and *z* describe the variation over the excitation volume. The presented simulation results correspond to a moderate powering of 1 mA. Fields for different powering values can be easily extrapolated from these profiles using a linear scaling. The magnetic fields generated by 1 mA input current in the proximity (within 5 μm) of the coil is about 5 Gauss, which is 1 order of magnitude higher than the Earth’s magnetic field; more turns of the coil would improve the strength and homogeneity of the magnetic field. However, the magnetic field decays quickly with distance from the coil, indicating that the generated magnetic field is highly localized and can be used to study single cells^2^. With improved fabrication design (for example, increasing the number of turns or the thickness of the coil), it is possible to enhance the generated magnetic field by 1 to 2 orders of magnitude, thus greatly further improving the sensitivity and performance of the microcoil-enabled cantilever.

**Figure S2.**
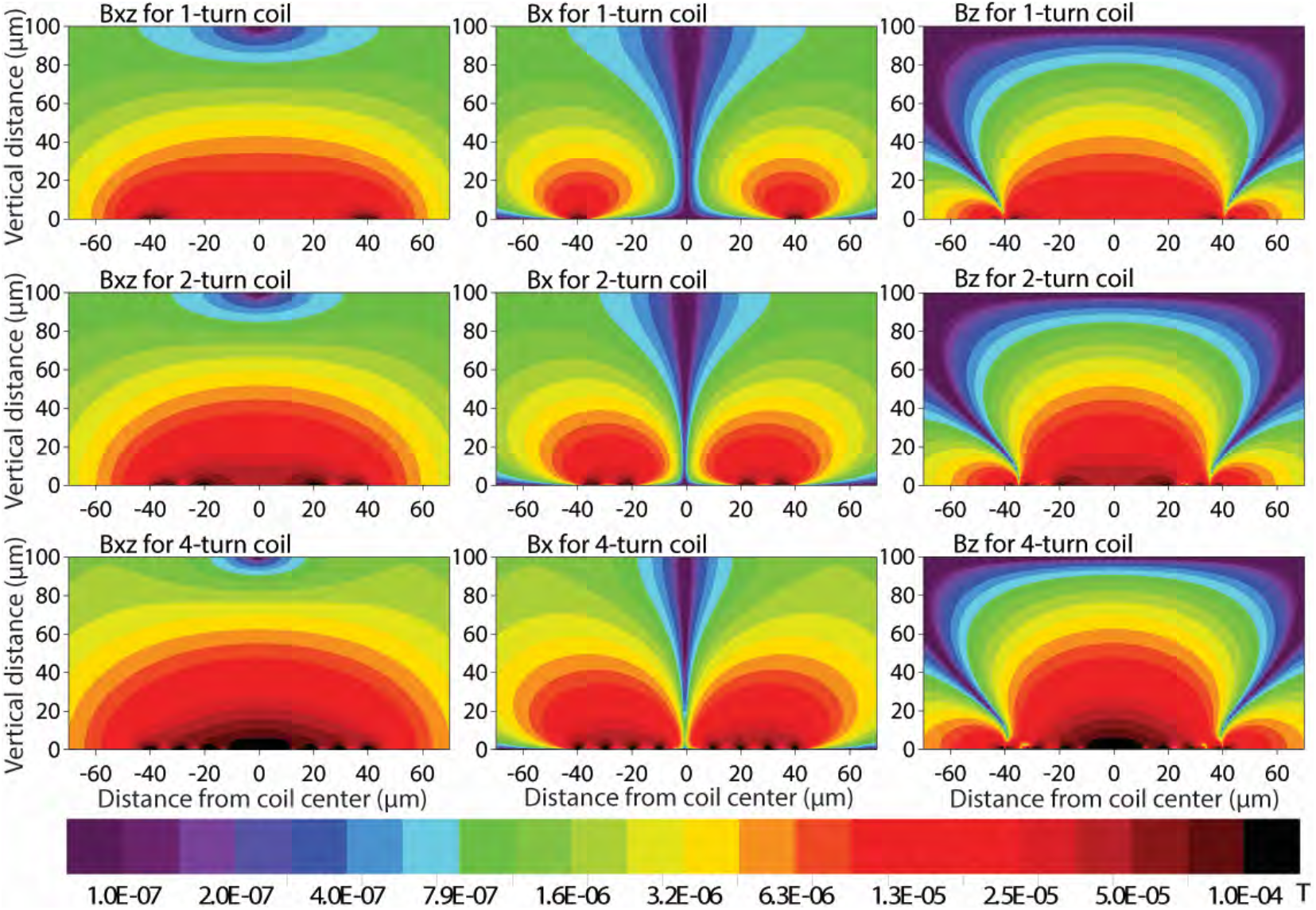
Finite element simulations of generated fields for 1 mA input current. Contour plots of the magnetic flux densities at planes vertical to the coil with span up to 70 um radial and 100 um vertical (coil center at x=0, y=0). Rows 1, 2, and 3 correspond to coils of 1, 2, and 4 turns respectively. Columns 1, 2, and 3 correspond to Bxz (vertical to the coil plane), Bx (along the cantilever width direction in the coil plane), and Bz (along the cantilever length direction in the coil plane), respectively.

### 4. Integration of Hybrid Coils and AFM System

The characterization and performance evaluation of the probes was conducted through a custom-modified KeySight 5500 AFM system (Keysight Technologies Inc., Santa Rosa, CA, USA). The system underwent two major modifications. First, a customized cantilever holder module was fabricated to attain both electrical and mechanical coupling to the hybrid probe (Figure S3). The second modification was the development of an aluminum AFM head holder that can be mounted on a wide-field fluorescence microscope. This was essential because it enabled simultaneous AFM scanning and fluorescence imaging for the physiological and chemical studies of cells, tissues, and biological samples. In addition, it replaced the commercial auto-approach sample stage that used magnets to secure the sample plate, thus prevented these magnets from inducing forces to the inductive path of the cantilever.

**Figure S3.**
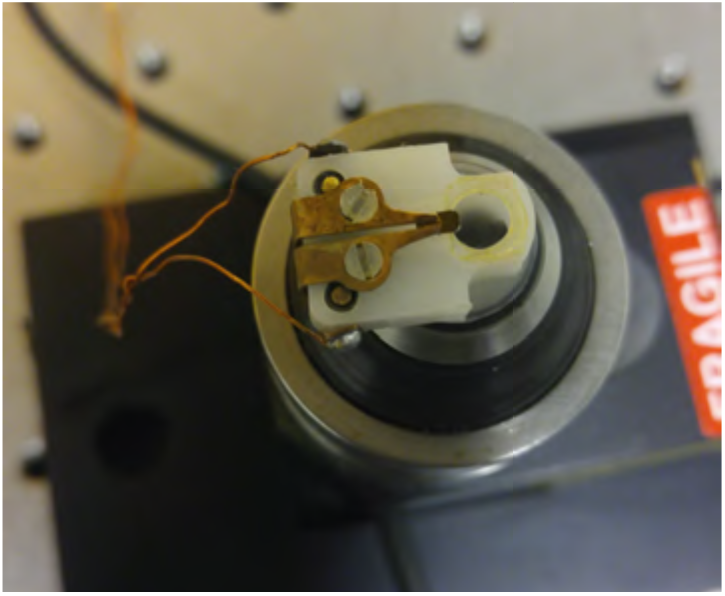
A custom-made cantilever holder module for the hybrid probe. This module provides both electrical and mechanical coupling to the hybrid probe.

### 5. Measurement of Magnetic Fields, Deflection, and Heating

#### 5.1 Magnetic field measurement

We measured the generated magnetic field in the proximity of the cantilever on unreleased devices, i.e. isolated device dies, cut in dimensions that approximated the AFM probe, with the silicon substrate present under the cantilever. The purpose of the experimental setup with this form of the device was to acquire the magnetic field at a fixed distance canceling out the deflection due to substrate heating. A magnetometer consisting of a three-dimensional axis configuration of magnetoresistive sensors (HMR2300, Honeywell, Morristown, NJ) was used for the measurement of the magnetic field. We focused on the *xy*-plane measurement of the field (with reference to the coordinate system of Figure S4) due to magnetometer geometry limitations. In particular, the *y*-component (along the length of the probe) of the magnetic flux density was recorded, since the large distance between the magnetoresistive sensor arrays on the magnetometer (compared to the size of the coil) did not allow for the confident simultaneous measurement of all the field components. In addition, the one-component study was considered adequate, since, except in the region of the feed-through traces, the coil geometry exhibits a cylindrical symmetry.

Initially, the optimal positioning of the device at the proximity of the *y*-oriented sensor of the magnetometer was decided based on the magnitude of the observed signal. Then, with the device engaged at the top of the sensing chip of the magnetometer, the current was swept from 0 to 10 mA with a step of 0.1 mA every 10 seconds using the current source (Sourcemeter 2440, Keithley, Cleveland, OH). The induced magnetic flux density values were recorded through the raw data captured at the serial port (RS232) of a laptop. The magnetometer provided 20 values per second, thus each point shown represents their mean value (Figure S5).

#### 5.2 Finite element simulation

In order to predict the cantilever deflection and temperature versus input power, we employed a finite element simulation tool (COMSOL Multiphysics) and performed parametric sweep studies for the one-turn coil, coupling three solution domains; a) solid mechanics, for the displacement of the cantilever, b) heat transfer in solids, and, c) electric currents. The geometric model was defined by the whole probe (Figure S4). The probe was rotated such that the simulation follows the actual experimental setup, where the probe was flipped horizontally after the mounting on the cantilever holder. In that way, the surfaces affecting heat convection had the correct orientation. Simulations were performed for the deflections and heating of the structure due to input currents ranging from 0.1 mA to 1.1 mA with steps of 0.1 mA, and the result is given in Figure S5.

**Figure S4.**
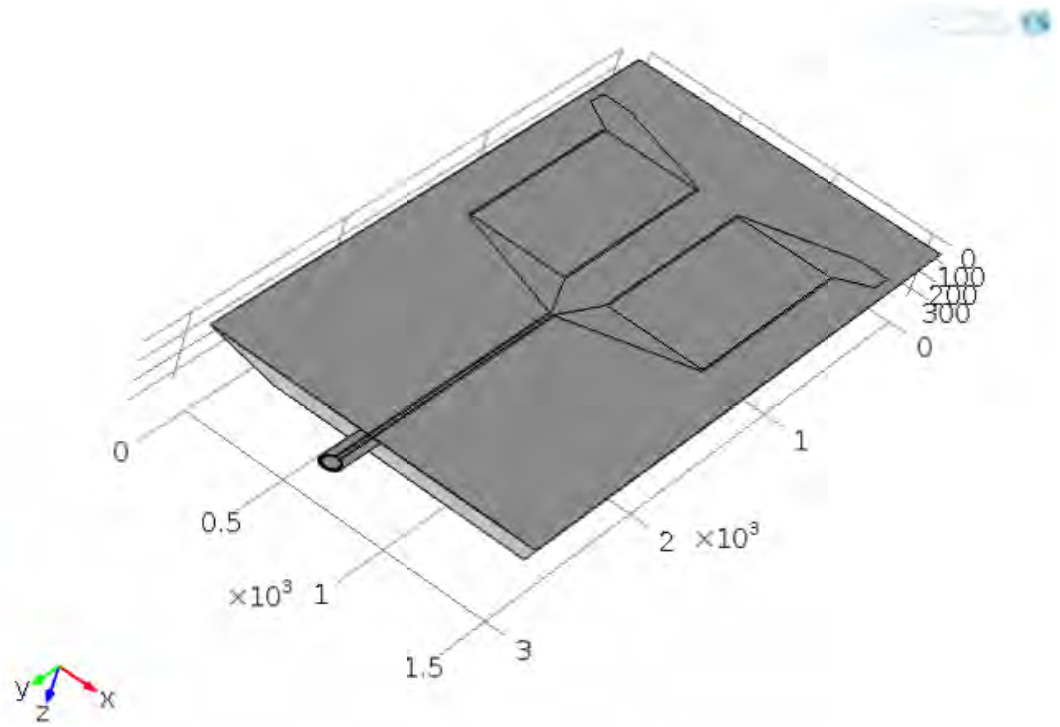
3D model of the one-turn cantilever for the finite element studies. The model was designed based on the actual dimensions of the probes after fabrication.

#### 5.3 Measurement of cantilever deflection due to Joule heating

We studied the Joule heating as a result of the current flow at the thin metallic traces of the cantilever. This investigation was necessary for two reasons. First, the heating of the cantilever induces cantilever deformation, expressed as vertical bending with reference to the direction normal to the plane of the coil. As a result, the magnetic flux density at a given point away from the cantilever decreased due to the induced deflection and increase of separation from the sample. Secondly, in magnetic force applications, the knowledge of the distance of the cantilever from the sample was important since the direction, magnitude, and gradient of the magnetic flux density needed to be monitored. For single-cell analysis, the deformation versus input power needs to be considered, since the distance between the coil and the cell has to remain constant or at the worst case fluctuate only at a small fraction, i.e. less than 1 micron, of their separation defined by the size of the attached tip, which is typically a sphere with diameter ranging from 5 to 30 microns.

To compare with our empirical observations, we employed a finite element simulation tool (COMSOL Multiphysics, see details in Supplementary Section 5.2) and performed parametric sweep studies for the 1-turn coil. Simulations were performed for the deflections and heating of the structure due to input currents ranging from 0.1 mA to 1.1 mA with steps of 0.1 mA. After applying a correction scaling factor (equal to 2.76), a very good agreement to the measured values was observed (Figure S5) for the deflection at the center of the coil. The use of a correction factor was necessary due to the difficulty in determining exact parameters of the heat transfer problem, e.g., the heat transfer coefficient h that was assumed to have the value of 5 W/(m2·K). Furthermore, the exact position of the laser spot on the cantilever greatly affects the measurement and, in practice, it is not easily controlled and reproduced. However, the presented model can provide intuition for the approximate position of the cantilever. The temperature at the center of the coil was also simulated and is given in Figure S5 after the application of the correction factor. Note that the coil was deposited on the top side of the cantilever, and the sample was at least 25 μm away from the coil with the cantilever and sphere tip in between. The simulations confirm that the cantilever temperature and deflection increase with input current; it is concluded, however, that the changes in temperature and deflection do not affect AFM operation when the input current is < 1 mA.

**Figure S5.**
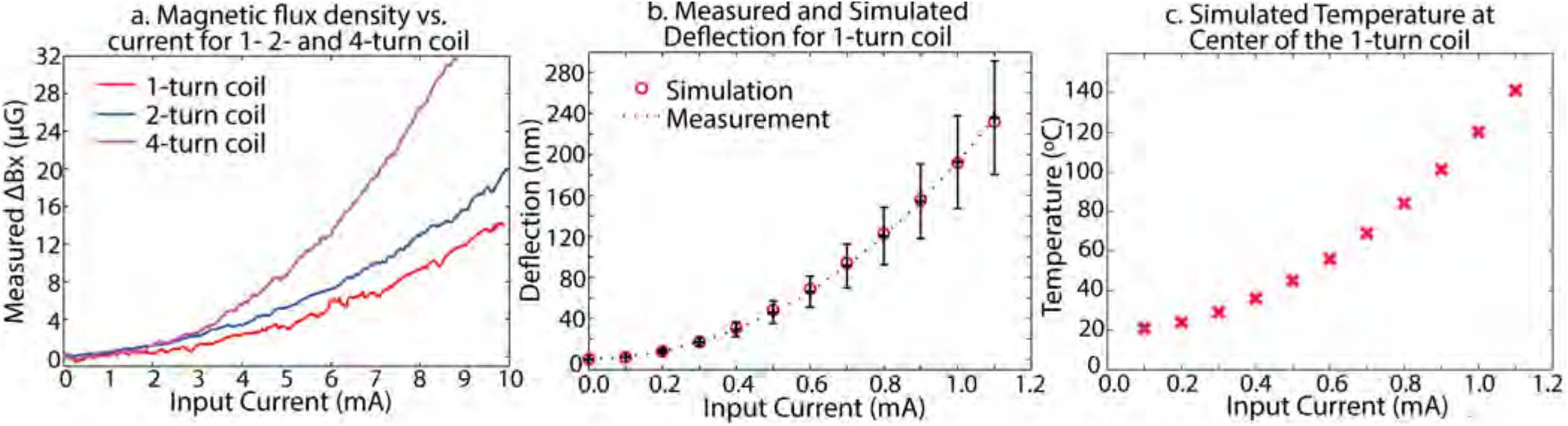
Magnetic field from the microcoils increases with input current at the cost of increased cantilever deflection and temperature. (a) Measured magnetic flux density variation vs. input current. (b) Cantilever deflection due to Joule heating. The error bars represent the standard deviation for a total number of 3 experiments. The large deviation, especially noticeable at high currents, might be due to the different location of laser spot on cantilevers between experiments. The presented finite element analysis results are for the point at the center of the coil. The agreement to measured values of deflection after the application of a correction factor of 2.76 is very good. The temperature at the same point is given in (c) simulated temperature at center of the 1-turn coil. The starting (room) temperature was set to be 20°C.

### 6. Hybrid probe for AFM techniques

A customized cantilever holder module was fabricated to attain both electrical and mechanical coupling to the hybrid probe. A MAC Mode III module (Keysight Technologies Inc., Santa Rosa, CA, USA) was used to tune the cantilever and supply AC current to the coil probe. Also a customized adapter was made for the Keysight 5500 AFM scanner to sit on top of a wide-field epifluorescent microscope (Eclipse Ti, Nikon Instruments Inc., Melville, NY, USA), allowing for simultaneous AFM scanning and optical microscopy.

#### 6.1 Frequency tuning in liquid

A one-turn coil probe was excited by AC current and dither piezo respectively in DI water; the thermal noise spectrum was also acquired (Figure S6). The amplitude was normalized with respect to the maximum value correspondingly. We found that the piezo drive generates many artificial resonant peaks, which are not necessarily related to the true cantilever resonances. In contrast, the current drive produces clean cantilever resonance response (confirmed by thermal spectrum), which is essential for quantitative dynamic mode AFM spectroscopy in liquid.

**Figure S6.**
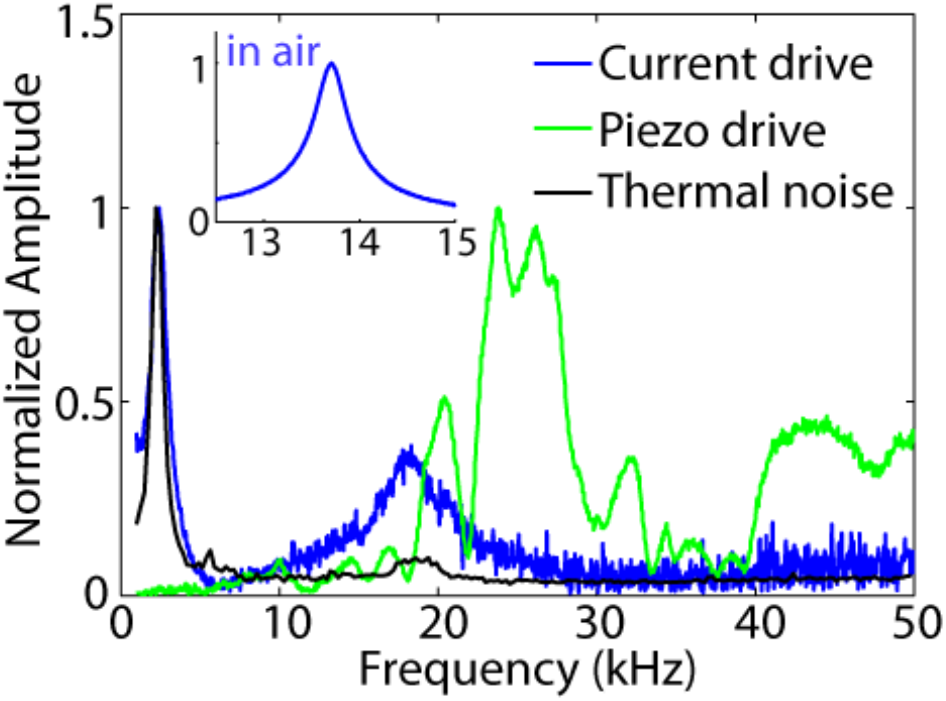
Frequency tuning curves of coil probe in liquid by different excitation mechanisms. The current drive produces clean cantilever resonance response even in liquid.

#### 6.2 Vibration amplitude enhancement by a small magnet

The flexural (bending) vibration amplitude of the coil probe can be greatly enhanced by a magnet placed under the coil. The 1^st^ bending eigenmode vibration amplitude as a function of gap between the coil and a ¼×¼×⅛” magnet at two different positions indicates that the stronger the magnetic field, the greater the vibration amplitude (Figure S7). Position 1 is at the center of the magnet, and position 2 is in between the center and the edge of the magnet.

**Figure S7.**
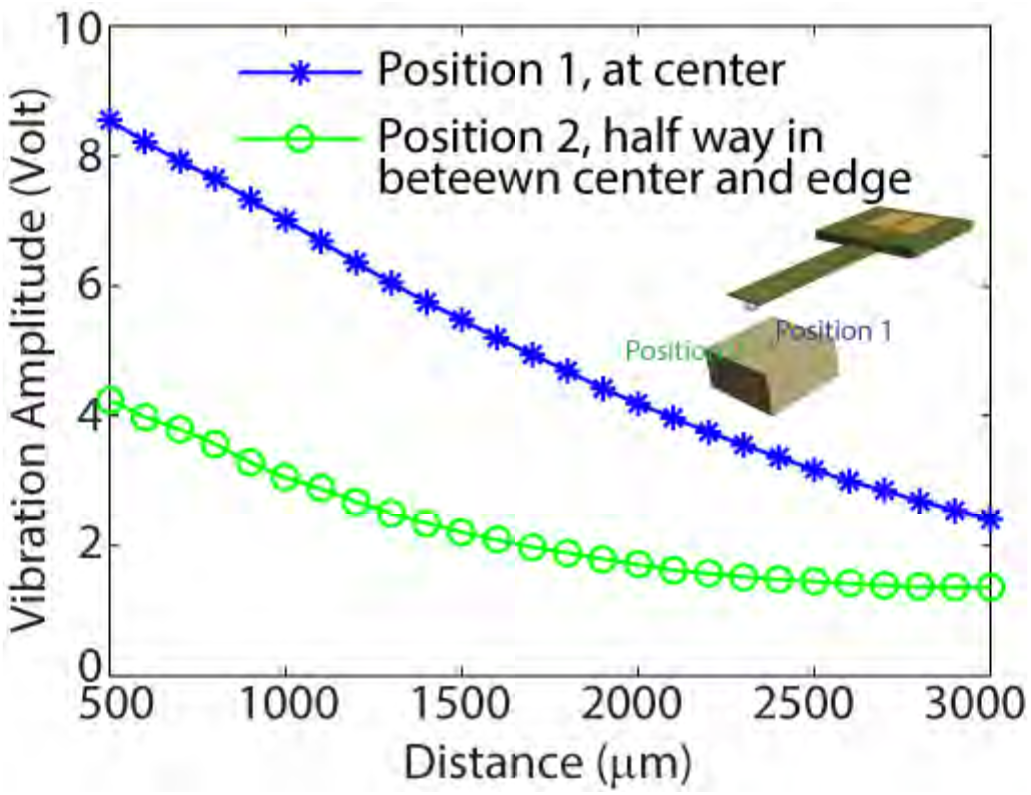
1^st^ bending eigenmode vibration amplitude increased as the gap between coil probe and magnet decreased at two different locations. At the center of the magnet (position 1), the magnetic field is strongest thus the enhancement of the bending vibration is greatest. Away from the center (position 2), the enhancement is roughly half as at the center. For both locations, the enhancement reduces as the vertical distance between the coil and the magnet increases.

#### 6.3 AFM imaging and sample preparation of PDMS embedded microbeads

To test the sensitivity for the stiffness and friction mapping technique by our hybrid probe, we first scanned a PDMS embedded microbeads sample. The original stock of 10 μm polystyrene (PS) microspheres (3.6 × 10^6^ beads⁄mL, Life Technologies, Grand Island, NY, USA) was diluted 100 times. A small 10 μL volume of the dilution was deposited on a mica surface (SPI Supplies, West Chester, PA) and dried by an air gun. Elastomeric polydimethylsiloxane (PDMS; Sylgard 184 Silicone Elastomer, Dow Corning, Auburn, MI, USA) was then deposited on the mica and cured at 60°C over night. When the PDMS was peeled off the mica, most microspheres transferred and were embedded on the PDMS surface (200-300 nm above the PDMS surface), and small portion of the surface was covered by thin layer of mica.

**Figure S8.**
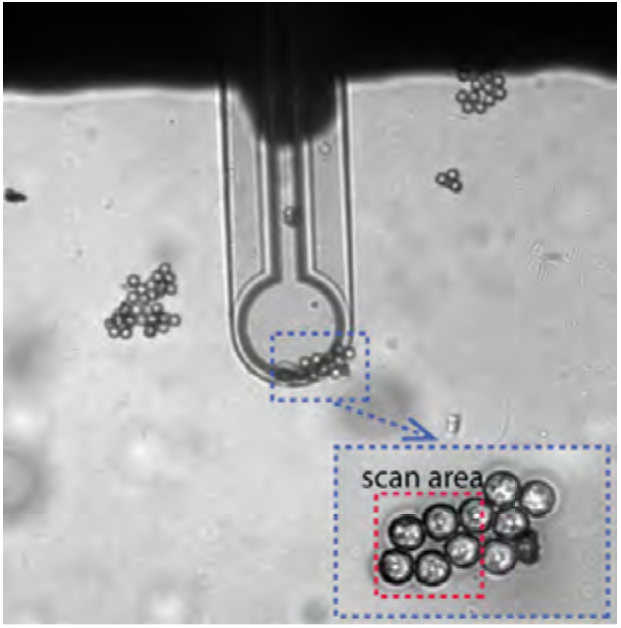
Coil probe is placed on top of a cluster of microbeads embedded in PDMS. With the help of an inverted optical microscope, the sample surface was clearly seen and the coil probe was easily positioned to the location desired.

#### 6.4 AFM imaging and sample preparation of PS-b-PMMA copolymer

To further test the sensitivity of our technique, we scanned a 1μm × 1μm small area on the surface of a ~50 nm thick film of poly(styrene-block-methyl methacrylate) (PS-b-PMMA). The two components of the copolymer have similar stiffness (bulk elastic moduli of PS and PMMA are 3.0 and 3.3 GPa, respectively^3^), however our technique successfully resolved the small variations (Figure 4(b)). Figure S9 shows that the lamellar nanostructures self-assembled by the PS-b-PMMA diblock copolymer in thin films (47 kg/mol PS and 53 kg/mol PMMA with a polydispersity index of 1.12, used as received from Polymer Source, Inc., Dorval, Quebec, Canada) had a periodicity of 47.5 nm.^4^

**Figure S9.**
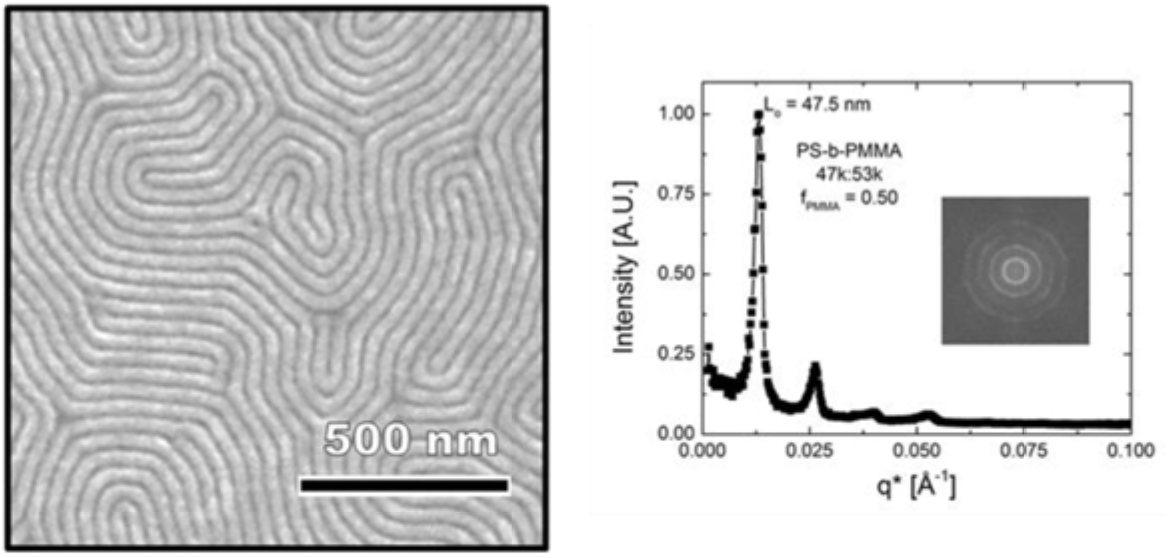
A scanning electron microscopy image (left) and fast Fourier transform (right) of lamellar nanostructures self-assembled in a PS-b-PMMA thin film. The 50 nm thick film had a lamellar morphology with a periodicity of 47.5 nm. The scale bar corresponds to 500 nm.

The process used to achieve lamellar nanostructures oriented perpendicular to the silicon substrate involved first pretreating the substrate with a crosslinking mat of random copolymer (poly(styrene-random-methyl methacrylate-random-glycidyl methacrylate) with 58 mol% PS synthesized by free-radical polymerization in-house) to create a neutral wetting surface. Briefly, a uniform film of PS-r-PMMA-r-PGMA was deposited by spin casting for a 0.3 wt% solution in anhydrous toluene on a cleaned Si wafer and baked for 4 h at 190 °C. Excess random copolymer that was not cross-linked to the substrate was subsequently removed by sonication in toluene. The block copolymer film was then spun cast from anhydrous toluene and solvent annealed in a saturated vapor environment of acetone and cyclohexane for 4 h to facilitate self-assembly into the lamellar domains. Solvent annealing was rapidly quenched by exposing the samples to air, and residual solvent in the film was removed by baking the sample for 30 min at 90 °C under a ~2 Torr vacuum.

The same sample was also scanned by a conventional probe in force volume mode for the purpose of comparison (Figure S10).

**Figure S10.**
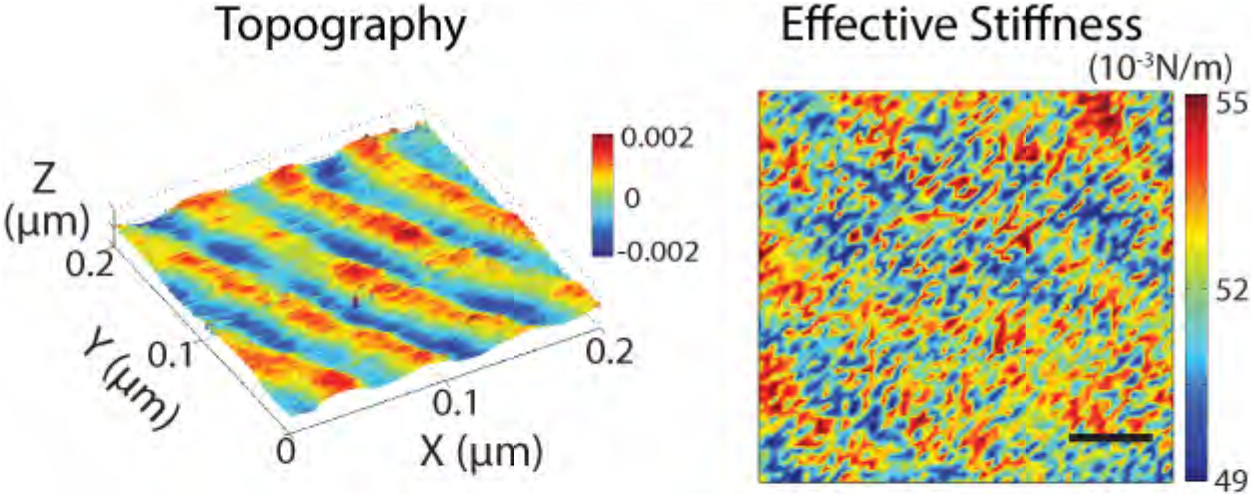
Force volume imaging of PS-b-PMMA copolymer with a regular AFM probe. Left: topography, right: effective stiffness map, scale bar: 40nm.

#### 6.5 AFM imaging and sample preparation of PMMA-cartilage interface

We also tested an osteochondral (articular cartilage-bone) sample harvested from the femoral head of a skeletally mature, male New Zealand white rabbit (Figure S11). The osteochondral sample was histologically dehydrated in a series of ethanol solutions, cleared in acetone, and embedded in poly(methyl methacrylate) (PMMA). The purpose of PMMA embedding was to ensure that the sample, containing an interface between the soft cartilage and stiff bone, could be easily micromilled to an optically flat finish. Raman spectroscopy (Renishaw Invia, 785 nm laser) was used to map peaks indicative of PMMA (810 cm^−1^).

**Figure S11.**
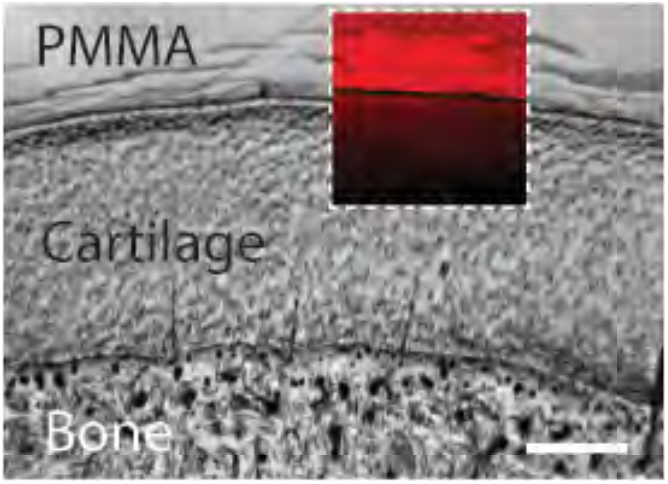
Wide field and Raman microscopy of PMMA-cartilage sample. Inset panel shows a map of the Raman intensity at 810 cm^−1^ corresponding to PMMA prevalence, artificially colored red. Image obtained using a 758 nm laser.

#### 6.6 Dynamic friction mapping

We also tested all the samples shown in Figure 4 for dynamic friction mapping (lateral force modulation imaging, Figure S12). For the PDMS sample covered by a thin layer of mica with an embedded microbead underneath, both piezo and current drive excitation methods showed little contrast except at the edges of the mica pieces (Figure S12(a)), indicating that the entire scan area was covered by mica. It is important to note that the stripe-like pattern on the effective friction map by coil probe was due to the laser reflection from the sample surface through the semi-transparent silicon nitride cantilever to the photo diode detector. This artifact can be greatly reduced if the sample surface is fabricated to be less reflective, like the PDMS and PS surfaces. In the future, the top side of the coil probe may be coated with a thin layer of aluminum to block the reflection from sample. For the PS-b-PMMA copolymer sample, since the sample surface is very flat, both excitation methods showed little contrast and the lamellar pattern is barely seen. For the cartilage-PMMA interface sample, due to the polishing processing, the cartilage and PMMA regions have similar roughness thus both excitation methods showed little contrast. However in all three cases, the torsional vibration amplitude of the current driven coil probe is significantly higher than the piezo driven regular probe even though the current drive uses a much smaller driving voltage (0.08 V) than the piezo drive (8 V). The torsional vibrations are very difficult to be excited off resonance with piezo excitation, our technique provides an efficient torsional actuation method for dynamic friction measurements and potentially can be applied in lateral contact stiffness measurements.

**Figure S12.**
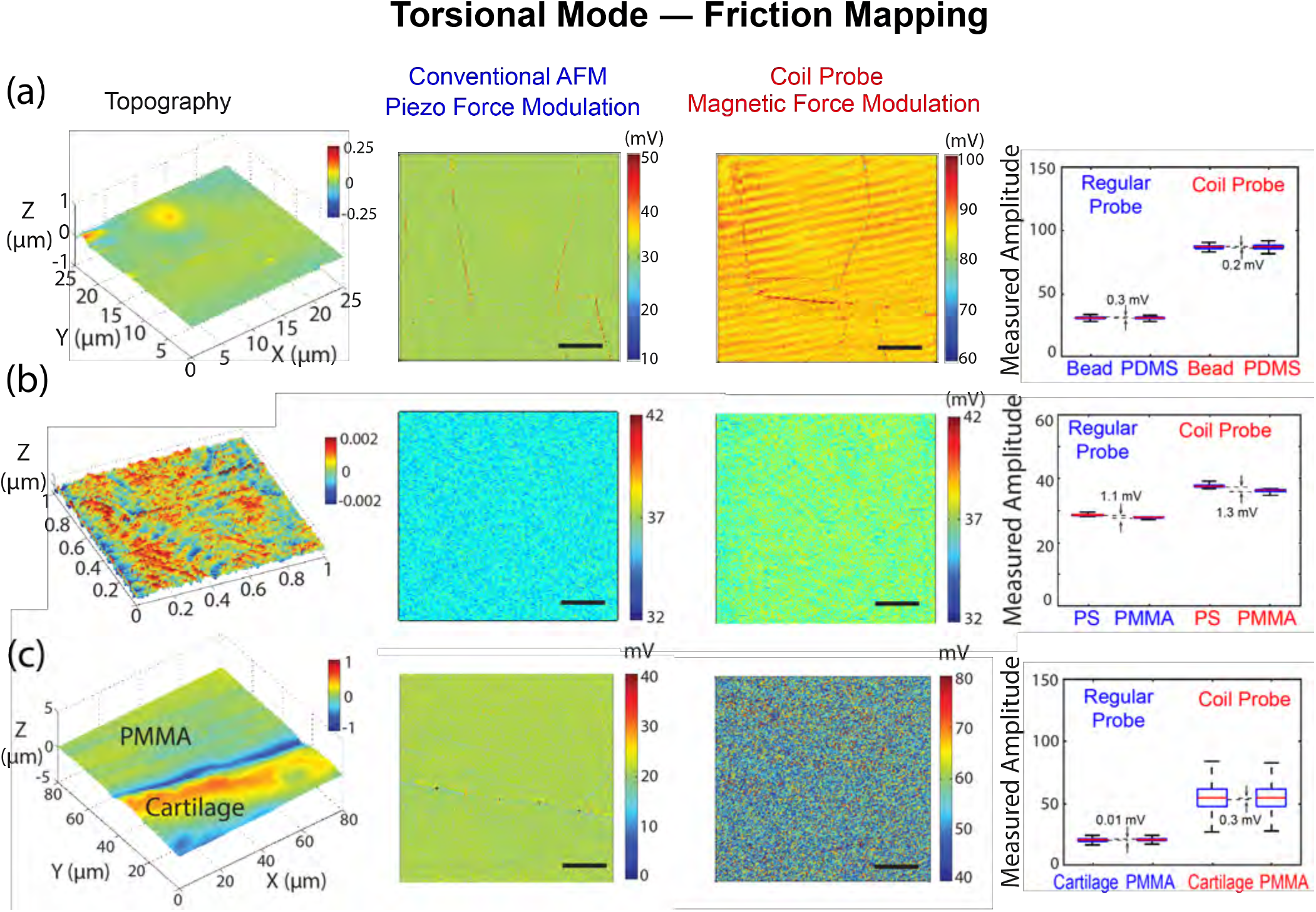
Dynamic friction mapping by torsional mode force modulation AFM. (a) A microbead embedded in PDMS covered by a thin layer of mica was scanned by regular probe with piezo drive and coil probe with current drive respectively, scale bars: 5 μm. (b) Surface of a thin film of PS-b-PMMA copolymer scanned by coil probe with piezo drive and current drive respectively, scale bars: 40 nm. (c) PMMA-cartilage sample was scanned by regular probe with piezo drive and coil probe with current drive respectively, scale bars: 20 μm.

### 7. Cell culture and microparticle injection

Primary Chondrocytes were harvested from the lateral and medial femoral condyles of five- to seven-month-old bovine knees. Cells were cultured (37°C, 5% CO_2_) in Dulbecco’s modified Eagle’s medium (DMEM/F12) supplemented with 10% fetal bovine serum (FBS), 0.1% bovine serum albumin, 50 ug/ml L-Ascorbate-2-phosphate, 100 units/mL penicillin, and 100 ug/mL streptomycin (Life Technologies, Carlsbad, CA).

Fluorescent superparamagnetic beads (0.9*μ*m; excitation/emission: 480nm/520nm; Bangs Labs, Fishers, IN) and carboxylate beads (1.1*μ*m; excitation/emission: 580nm/605nm; Life Technologies) were prepared (at 120*μ*g/ml) in DI water. Two 35mm petri dishes were plated with 1.5 million cells each and allowed to sit for no less than 3 hours. The media was then removed, and the microparticles were ballistically injected (35% injection rate; 1350 psi; Biolistic PDS-1000/HE, BioRad, Richmond, CA) into the cytoplasm or nucleus of the cells (Figure 5(a-b)). The cells were incubated for 8-24 hours, released (TrypLE Express™; Life Technologies) and suspended together, and then replated on 35mm Cellview™ cell culture dishes (Greiner Bio-One, Frickenhausen, Germany). After a 1 hour equilibration, chondrocytes were washed with medium and stained with Hoechst and Calcein AM (Life Technologies) prior to imaging. Confocal microscopy (Figure 5; AIR_MP, Nikon) confirmed the penetration of the microbeads into the cytoplasm and nuclei.

### 8. Magnetic manipulation testing and imaging

Microbead movements were accomplished using a 2- or 4-turn microcoil and custom AFM nosecone that was immersed in media and lowered to within 10-30 um of the chondrocyte monolayer. A current pulse (0.2mA, 0.5s, 0.1 Hz) was passed to the cantilever by a function waveform generator (Agilent). Cells were simultaneously imaged using widefield microscopy (Eclipse Ti, Nikon) and a CCD camera (iXon+, Andor, Belfast, NIR) for 60s undisturbed, 120s with pulsing, and a final 120s minutes undisturbed, at high spatiotemporal resolution (180nm/px, 1.2s frame rate). Microbead positions were analyzed using semiautomated threshold and particle centroid-tracking routines (ImageJ and MATLAB The Mathworks, Natick, MA) with a 15nm spatial resolution^5^.

The forces applied to the superparamagnetic beads were estimated using the standard formula for weak magnetic fields^6^:

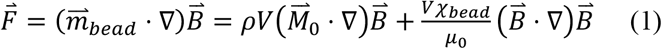

where *V* is the volume of the magnetic beads (3.82e-19m^3^), *χ*_*bead*_ is the magnetic susceptibility of the beads, (0.17, supplied by manufacturer), *μ*_0_ = 4*π* × 10^−7^(*TmA*^−1^), the permeability of a vacuum. With a current of 0.2mA, 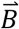 at 10-30 μm is 1.3e-5 T, resulting in femtonewton (fN) forces.

Post testing, particle movements were analyzed. The particle mean standard displacements (MSDs) were equated overtime (Figure 5(a)). The MSDs were binned into 30s increments yielding average centroid movements. The MSDs were then offset to the first 30s of Brownian motion. (Figure 5, *n*=27). Data was analyzed by means of a two-way ANOVA. Magnetic property and time were treated as categorical data with mean displacement of the microparticles as the dependent variable. Magnetic field application is embedded in the time course series. Results were found to be significant if p<0.05.

